# α-Synuclein structural heterogeneity associates with distinct cytopathologies in multiple system atrophy

**DOI:** 10.64898/2026.11.13.766536

**Authors:** Xiaoxiao Xu, Shelley L. Forrest, Ivan Martinez-Valbuena, Carly Scramstad, Ekaterina Rogaeva, Suzanne E. Kosteniuk, Joel C. Watts, Gabor G. Kovacs, Masahiro Enomoto

## Abstract

Multiple system atrophy (MSA) is characterized by oligodendroglial α-synuclein inclusions, although neuronal inclusions also occur. We investigated whether regional differences in α- synuclein filament structure are associated with distinct cytopathologies in a neuropathologically confirmed MSA brain by comparing the medial temporal lobe, which showed prominent neuronal inclusions and limited oligodendroglial pathology, with cerebellar white matter, which contained exclusively oligodendroglial inclusions. Cryo-electron microscopy resolved filaments from these regions at 3.0 Å and 2.0 Å, respectively. Medial temporal filaments contained a previously undescribed protofilament fold, PF-IIIA, paired with canonical PF-IIB2, whereas cerebellar filaments adopted the canonical MSA Type II structure. PF-IIIA exhibits extensive remodeling of its core while retaining the asymmetric doublet architecture. These findings reveal regional α- synuclein structural heterogeneity within a single MSA brain and suggest a potential association between distinct filament conformations and neuronal versus oligodendroglial cytopathology, highlighting the importance of regional and cellular context in filament organization and disease heterogeneity.

## Introduction

Multiple system atrophy (MSA) is a progressive neurodegenerative disease characterized by α- synuclein filament accumulation, predominantly within oligodendroglial cytoplasmic inclusions^1^. Together with Parkinson’s disease and dementia with Lewy bodies, MSA constitutes one of the major synucleinopathies^2,3^. Its neuropathology is characterized by striatonigral and olivopontocerebellar degeneration, which broadly correspond to the parkinsonian and cerebellar clinical phenotypes, respectively^1^. However, α-synuclein pathology is not restricted to oligodendrocytes; neuronal cytoplasmic inclusions also occur and exhibit ultrastructural heterogeneity, including fibrillar and membranous forms resembling those observed in Parkinson’s disease^4^. The coexistence of neuronal and oligodendroglial inclusions raises the question of whether these inclusion types contain structurally distinct α-synuclein filaments.

Cryo-electron microscopy (cryo-EM) has provided a structural basis for distinguishing α- synuclein assemblies associated with different synucleinopathies. Brain-derived MSA filaments comprise two major types, each assembled from two structurally distinct protofilaments, designated PF-A and PF-B^5,6^. These asymmetric doublet assemblies differ from the single- protofilament Lewy fold identified in Parkinson’s disease and dementia with Lewy bodies^7^. Subsequent work identified a low-abundance MSA filament subtype, Type I2, in one individual, containing a chimeric protofilament with structural features of both Type I and Type II folds^8^. We previously identified a Lewy–MSA hybrid fold in an atypical MSA case, combining structural features of the Lewy and MSA folds^4^. Together, these findings expand the known spectrum of α- synuclein conformations and raise the question of how this diversity relates to MSA cytopathology.

The anatomical and cellular distribution of these α-synuclein folds remains unresolved. Although the relative abundance of MSA filament types varies among individuals, its relationship to clinical phenotypes and regional cytopathology remains unclear^4,6,8,9^. In particular, whether neuronal and oligodendroglial inclusions contain distinct filament folds remains unsolved^4,10^. Comparing regions with contrasting neuronal and oligodendroglial involvement allows filament structures to be examined alongside the inclusion types present in the sampled tissue. Comparisons within a single individual also avoid genetic and clinical differences between patients.

MSA cases with prominent limbic neuronal inclusions, known as atypical MSA, are particularly informative in this regard. Core MSA-affected regions, such as the striatum and pons, contain both neuronal and oligodendroglial α-synuclein pathology^11,12^, making it difficult to associate folds recovered from tissue homogenates with either cell type. In contrast, comparing the medial temporal lobe, which shows more prominent neuronal pathology, with cerebellar white matter, which shows predominantly oligodendroglial pathology, allows regional filament structures to be compared in the context of differing cellular pathology, although it does not directly assign folds to either cell type. Such analyses can also establish whether distinct folds coexist within a single brain. This approach complements comparisons across diagnostic groups by examining structural heterogeneity within a single neuropathologically defined disease.

Here, we report a previously undescribed α-synuclein protofilament fold in a neuropathologically confirmed MSA case with prominent limbic neuronal inclusions. Cryo-EM analysis of sarkosyl-insoluble filaments from the medial temporal lobe revealed a novel PF-A fold, designated PF-IIIA, paired with canonical PF-IIB2. PF-IIIA exhibits extensive remodeling of the protofilament core while preserving the overall MSA Type II doublet architecture. In contrast, cerebellar filaments isolated from the same individual adopt the canonical MSA Type II structure. These findings expand the known structural diversity of MSA-associated α-synuclein filaments and demonstrate that distinct PF-A folds can coexist across different anatomical regions within a single brain.

## Results Case Report

A 65-year-old man presented with a progressive movement disorder with a total disease duration of approximately 12 years, from symptom onset to death. Initial symptoms comprised left arm tremor, generalized bradykinesia, and gait difficulty. On examination approximately one year after onset, the tremor abated with intention, and he demonstrated increased tone, bradykinesia, an unsteady gait, mild saccadic pursuit, hypophonia, and hypomimia. Early non-motor features included orthostatic hypotension, constipation, bladder dysfunction, daytime somnolence, and depression/anxiety. Over the disease course he developed additional features including hypotension, anterocollis, dysphagia, palilalia, lateral (right-sided) truncal leaning, and weight loss. A psychiatric consultation approximately 10 years after onset noted perseverations possibly attributable to cognitive change or anxiety, and notes in the final year of illness described mild distractibility. A pre-existing deficit in cognitive processing speed was attributed to a traumatic brain injury sustained approximately two years before onset. MRI images and reports were not available in the reviewed records. The patient had a mild response to pramipexole and a modest response to oral levodopa with rapid wearing-off, but a good response to intrajejunal levodopa– carbidopa. There was no documented family history of movement disorders. The clinical diagnosis was recorded as Parkinson’s disease versus possible MSA. The patient died approximately 12 years after symptom onset. His medical history was notable for a traumatic brain injury (following a fall on ice) sustained approximately two years before onset, with basal skull fracture and craniotomy, resulting in language, cognitive, and olfactory changes; furthermore, hypothyroidism and nephrolithiasis were also documented.

## Neuropathological and genetic findings

The right hemisphere was formalin-fixed and the left was frozen. Macroscopic examination showed thickened leptomeninges with focal subarachnoid blood over the right inferior parietal region, but an unremarkable gyral pattern without appreciable cortical or mesial temporal atrophy. The ventricles were only mildly widened. The putamen showed brownish discoloration and marked shrinkage. Brainstem sections revealed moderate depigmentation of the substantia nigra and locus coeruleus, with a mildly reduced pontine base calibre; the cerebellar cortex showed mild atrophy.

Histologically, neuronal loss and gliosis were most severe in the putamen, moderate in the substantia nigra, and mild in the motor cortex, globus pallidus, subthalamic nucleus, locus coeruleus, pontine base, inferior olive, and cerebellum (Purkinje and granule cell layers). A single focal subarachnoid bleed was present in the right inferior parietal cortex. Arteriolosclerosis was prominent in the frontal and occipital cortex, hippocampus, and basal ganglia. Amyloid angiopathy, ballooned neurons, and inflammatory infiltrates were absent.

Immunohistochemistry for α-synuclein (Phosphorylated α-synuclein and the 5G4 antibody) on the FFPE samples (**Fig 1. A-H**) of the right hemisphere demonstrated widespread glial cytoplasmic inclusions (Papp–Lantos bodies) throughout the cortical and cerebellar white matter, basal ganglia, and brainstem, most abundant in the dorsolateral putamen, motor cortex, pontine base, and cerebellar white matter. Notably, α-synuclein–positive neuronal cytoplasmic inclusions, including spherical and ring-like forms, were present in moderate degree in the hippocampus (CA1 and dentate gyrus) and amygdala, with corresponding p62 immunoreactivity. Phosphorylated α- synuclein antibody confirmed these neuronal and oligodendroglial inclusions in frozen, post-fixed tissue of the left hemisphere (**Fig 1. I-P**).

**Figure 1.**
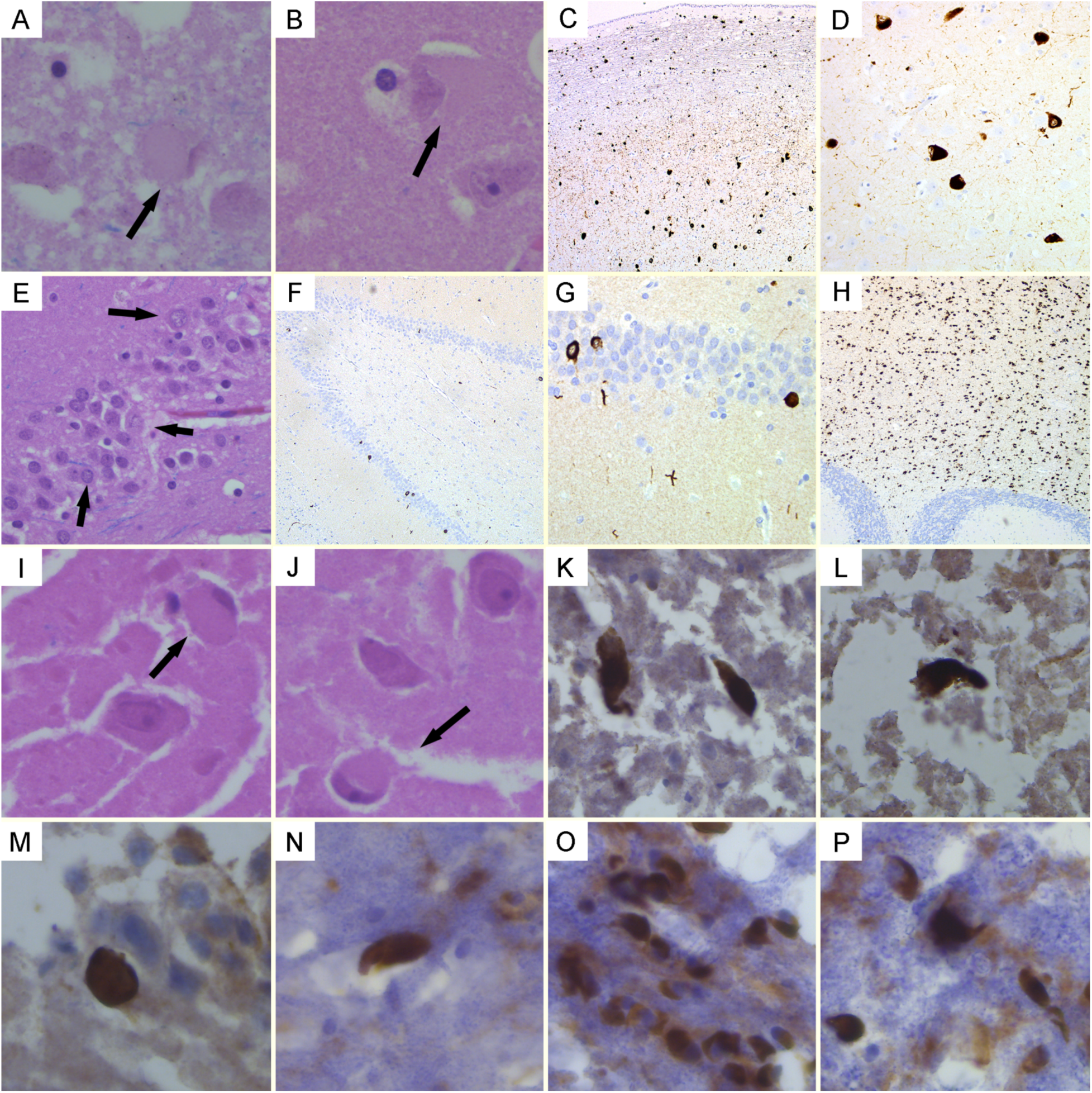
: (A–H) Formalin-fixed, paraffin-embedded (FFPE) tissue sections. (A, **B)** Hematoxylin–eosin (HE) staining, with (b) at higher magnification, showing pyramidal neurons of the CA1 region bearing eosinophilic round and horseshoe-shaped intracytoplasmic inclusions. **(C, D)** Immunostaining with the 5G4 anti–α-synuclein antibody: **(C)** at lower magnification, demonstrating oligodendroglial inclusions in the fimbria of the hippocampus together with neuronal cytoplasmic inclusions (lower part of the image); **(D)** higher magnification of CA1 pyramidal neurons with cytoplasmic inclusions. **(E)** HE staining of the dentate gyrus granule cell layer showing ring-like inclusions (arrows). **(F, G)** 5G4 α-synuclein immunostaining of the dentate gyrus at lower (**F**) and higher (**G**) magnification. **(H)** 5G4 α-synuclein immunostaining of the cerebellum revealing abundant oligodendroglial cytoplasmic inclusions. **(I–P) Frozen, post-fixed tissue sections. (I, J)** HE staining of the hippocampus revealing rounded eosinophilic inclusions (arrows). **(K)** Immunostaining for disease-associated α-synuclein revealing the neuronal inclusions, further confirmed by **(L)** p62 immunostaining. **(M- P)** Immunostaining for phosphorylated α-synuclein revealing neuronal inclusions in the granule cell layer of the dentate gyrus **(M)** and oligodendroglial cytoplasmic inclusions in the white matter of the frontal lobe **(N)**, the cerebellum **(O)**, and the striatum **(P)**.

Concomitant Alzheimer-type pathology was limited: tau immunostaining showed neurofibrillary tangles and neuropil threads restricted to the entorhinal cortex and hippocampus with a few amygdala tangles (Braak stage II), and amyloid-β revealed only sparse diffuse plaques (Thal phase 2) without cerebral amyloid angiopathy. p-TDP-43 was negative throughout.

In summary, the findings were diagnostic of MSA, distinguished by moderate degree limbic/hippocampal neuronal α-synuclein pathology. Coexistent low-level Alzheimer’s disease neuropathologic change (A1B1C0) was present, without significant TDP-43 pathology.

Genetic analysis using Nitrobacter array did not detect any known pathogenic mutations in genes linked to neurodegenerative diseases, including multiplications at the *SNCA* locus (**Fig. S1**).

## Cryo-EM structures of medial temporal and cerebellar α-synuclein filaments

To investigate whether the prominent limbic neuronal α-synuclein pathology was accompanied by regional differences in filament structure, we examined α-synuclein filaments extracted from the medial temporal lobe, including the hippocampus and amygdala, and from the cerebellum of the same individual by cryo-EM. The medial temporal region exhibited both neuronal and oligodendroglial α-synuclein inclusions in both hemispheres, whereas cerebellar pathology was characterized exclusively by oligodendroglial inclusions, with no neuronal cytoplasmic inclusions observed in the examined sections (**Fig. 1**). Filaments were extracted from 1–2 g of tissue per region and applied to graphene oxide–coated cryo-EM grids. This preparation yielded sufficient filaments for high-resolution structure determination from both regions. Based on 2D classification of helical segments, approximately 60% of classified segments from the medial temporal lobe and 75% from the cerebellum displayed a twisted morphology, enabling structure determination by helical reconstruction (**Fig. S2**). Reconstructions of the medial temporal lobe– and cerebellum- derived filaments (hereafter MT-filament and Cere-filament, respectively) were resolved to 3.0 Å and 2.0 Å, respectively (**Figs. 2A, S3, S4** **and Table S1**).

**Figure 2.**
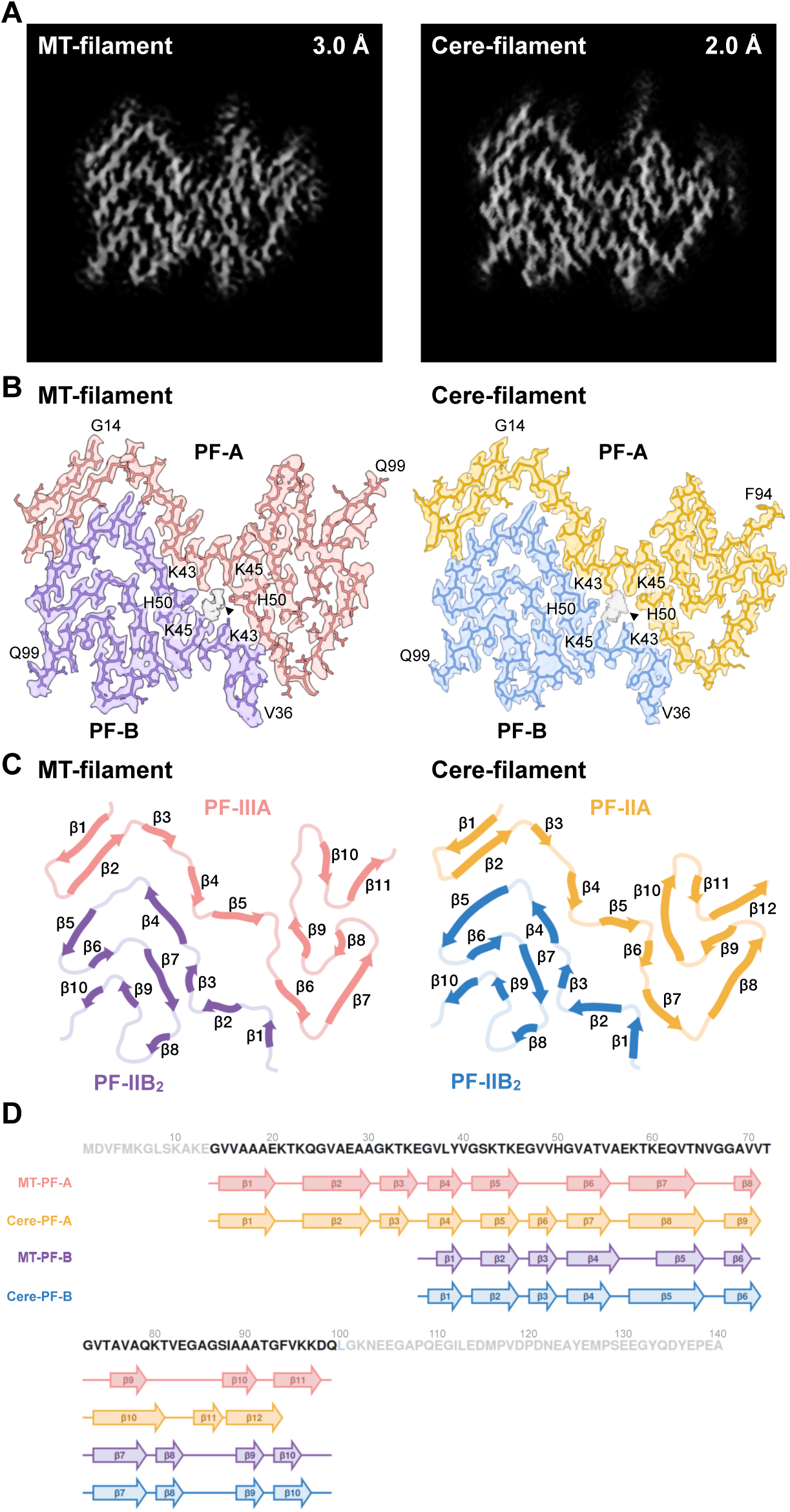
Cryo-EM maps and atomic models of a-synuclein filaments. **(A)** Cross-sections of cryo-EM maps of from the medial temporal lobe (MT-filament, left) and from the cerebellum (Cere-filament, right). **(B)** Sharpened high-resolution cryo-EM maps with overlaid atomic models of the MT-filament (left) and Cere-filament (right). Black arrowheads indicate non-proteinaceous density, shown in gray. **(C)** Schematic of secondary structure elements in the MT-filament (left) and Cere-filament (right). **(D)** Schematic of the human α-synuclein sequence, comparing secondary structure elements in the protofilament (PF) cores from the MT- and Cere-filaments. Residues resolved in at least one PF core are depicted in black.

**Figure 3.**
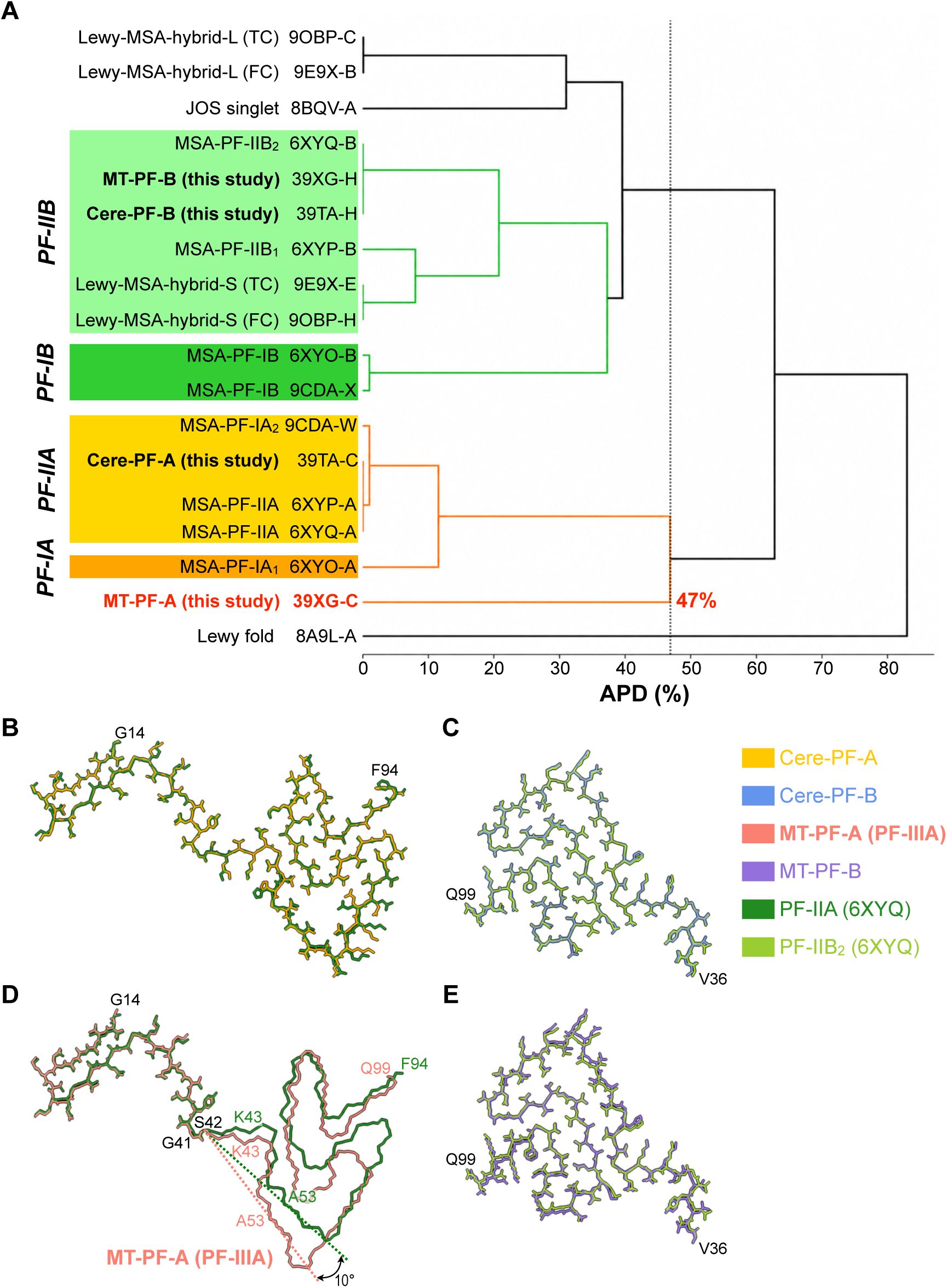
Structural comparison of MT- and Cere-filaments. **(A)** Clustering of α-synuclein protofilament folds based on pairwise APD values. PDB codes and chain IDs are listed as “PDB code-chain ID” for all clustered structures. Dendrogram branches are colored to highlight the PF-A and PF-B folds, and a vertical dotted line indicates the APD value of 47% separating PF-IIA and MT-PF-A (PF-IIIA). **(B-E)** Overlaid structures of Cere-PF-A, Cere- PF-B, MT-PF-A (PF-IIIA), or MT-PF-B with canonical PF-IIA or PF-IIB2 structures from PDB 6XYQ. Entire structures are superimposed in **(B, C, and E)**, whereas G14-G41 are superimposed in **(D)**. In **(D)**, only Cα backbone traces are shown from S42 onward to highlight difference between MT-PF-A (PF-IIIA) and PF-IIA. Dotted line highlight the ∼10° orientation difference of K43-A53 between PF-IIIA and PF-IIA after G14-G41 alignment.

**Figure 4.**
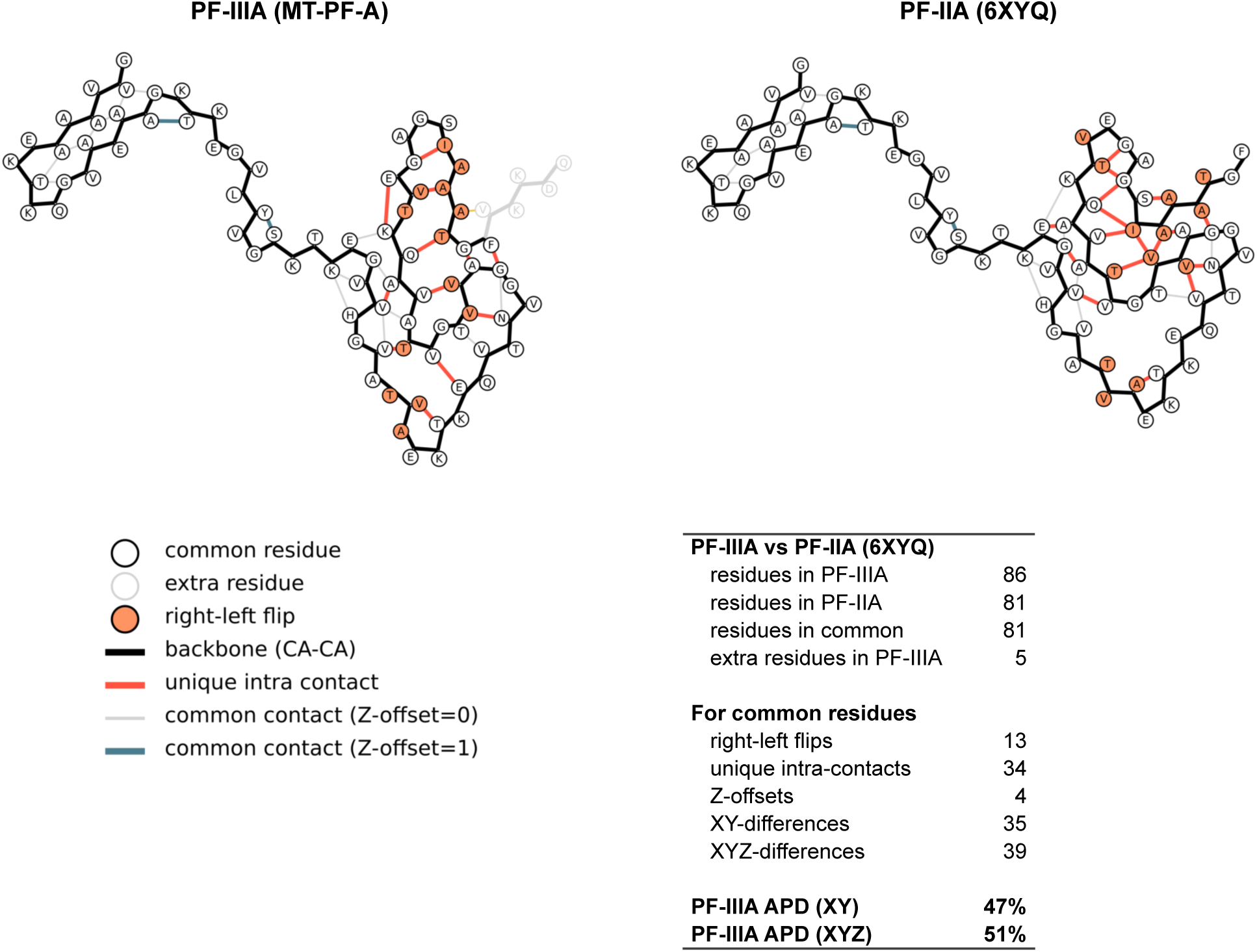
PF-IIIA exhibits distinct side-chain orientations and packing interactions. Amyloid packing difference (APD) schematic comparison of PF-IIIA from the medial temporal lobe (MT-PF-A; chain C; left) and canonical PF-IIA (PDB: 6XYQ; chain A; right). Circles indicate residues, and black lines trace the Cα backbone. White circles indicate residues present in both structures, pale gray circles indicate residues resolved in only one structure, and orange circles indicate side-chain orientation flips. Contacts are defined as a distance of ≤6.5 Å between non-hydrogen side-chain atoms (Cα for glycine), subject to sequence-separation and side-chain orientation criteria. Red lines indicate intra-protofilament contacts unique to either structure. Conserved contacts are colored by their Z-offset, calculated as the difference in Z coordinates of the contacting residues’ Cα atoms in the middle layer, divided by 4.75 Å and rounded to an integer. XY-APD excludes these axial offsets, whereas XYZ-APD includes them, yielding values of 47% and 51%, respectively.

Consistent with previously reported ex vivo α-synuclein filament structures from MSA patients^5,6,8,9^, both structures comprise doublet filaments in which two protofilaments (PF-A and PF-B) pack asymmetrically through an extended interface spanning nearly the entire width of the filament core (**Fig. 2B**). PF-A and PF-B of the MT-filament were resolved from G14 to Q99 and from V36 to Q99, respectively, whereas PF-A and PF-B of the Cere-filament were resolved from G14 to F94 and from V36 to Q99, respectively (**Fig. 2B**). In both structures, basic residues K43, K45, and H50 from each protofilament surround an unassigned density at the center of the filament core (**Fig. 2B**). Inter-protofilament packing is further stabilized by salt bridges between E28 of PF- A and K60 of PF-B and between E35 of PF-A and K58 of PF-B, together with close packing between A53-V55 of PF-A and the β1–β2 loop of PF-B (**Fig. S5**).

## Medial temporal filaments contain a novel PF-A fold

To compare the protofilament folds of the MT-filament and Cere-filament, we employed amyloid packing difference (APD)—a metric that expresses the percentage of residues differing in cross-β packing contacts or side-chain orientations, or ordered in only one of the two structures^13^— alongside Cα root-mean-square deviation (Cα RMSD) analysis. The two major previously reported MSA filament types, Type I and Type II, each comprise two distinct protofilaments, designated PF-IA/PF-IB and PF-IIA/PF-IIB, respectively^5,6^. Hierarchical clustering of α-synuclein protofilament folds based on all pairwise XY-APD values—a variant of APD that ignores whether cross-β packing interactions occur within the same molecular rung or across different rungs^13^— shows that PF-A and PF-B folds diverge at an XY-APD of exceeding 60%, whereas the Lewy fold separates from all other α-synuclein folds at XY-APD values exceeding 80% (**Fig. 3A**). At an XY- APD of 40%, the PF-B cluster containing canonical PF-IB and PF-IIB folds joins a cluster comprising the juvenile-onset synucleinopathy (JOS) fold^14^ and the Lewy–MSA hybrid fold identified in an atypical MSA (FTLD-synuclein) case^4^.

Cere-PF-A and Cere-PF-B exhibited XY-APD values of 0% relative to the previously reported PF-IIA (PDB 6XYP and 6XYQ) and PF-IIB2 (PDB 6XYQ), respectively, indicating conservation of the packing contacts and residue orientations assessed by this metric (**Fig. 3A**). Consistent with these assignments, superposition onto PF-IIA and PF-IIB2 from PDB 6XYQ yielded Cα RMSD values of 0.372 Å and 0.299 Å, respectively (**Figs. 3B and 3C**). We therefore designated the two protofilaments of the Cere-filament as PF-IIA and PF-IIB2 (**Figs. 2C and 2D**). Strikingly, MT-PF- A branched from the cluster comprising canonical PF-IA and PF-IIA folds at an XY-APD value of 47%, whereas MT-PF-B showed an XY-APD of 0% relative to PF-IIB2 (**Fig. 3A**). Consistent with this classification, MT-PF-B superimposed closely onto PF-IIB2 from PDB 6XYQ, yielding a Cα RMSD of 0.770 Å (**Fig. 3E**). These results demonstrate that the MT-filament comprises a novel PF-A fold, designated PF-IIIA, paired with a canonical PF-IIB2 protofilament (**Figs. 2C and 2D**).

## PF-IIIA exhibits extensive remodeling of the protofilament core

Structural comparison of PF-IIIA with the canonical PF-IIA fold (PDB 6XYQ) revealed that residues G14–G41 superimpose closely, with a Cα RMSD of 0.901 Å (**Fig. 3D**). The conformational difference includes an approximately 10° relative rotation between the G14–G41 and K43–A53 segments, with S42 at their junction (**Fig. 3D)**. In contrast, the T54–Q99 segment of PF-IIIA adopts a markedly different conformation from the corresponding T54–F94 segment of PF-IIA (**Figs. 3D**).

To characterize these differences, we used APD schematic representations to compare residue orientations and cross-β packing interactions with those of canonical PF-IIA (6XYQ). Cere-PF-A retained the same residue orientations, packing contacts, and axial offsets assessed by APD, yielding XY-APD and XYZ-APD values of 0% (**Fig. S5**). In contrast, PF-IIIA showed substantial differences in side-chain orientations and packing interactions, with XY- and XYZ-APD values of 47% and 51%, respectively (**Fig. 4**). Flips of the T54, V55, and A56 side chains accompany a PF-IIIA-specific packing interaction between V55 and T59, whereas a flip of the T75 side chain accompanies PF-IIIA-specific packing interactions between T75 and V52 and between E61 and V74. These changes form part of an extensive rearrangement of the packing network across the C- terminal portion of the resolved core. The side-chain rearrangements around T54–A56 are also accompanied by differences in inter-protofilament packing, with close contacts involving V52– G41 and A53–V40 in the Cere-filament, compared with more localized contacts involving T54– V40 and A53–V40 in the MT-filament (**Fig. S4**). Therefore, whereas Cere-PF-A closely preserves the canonical PF-IIA fold, PF-IIIA undergoes substantial remodeling of the downstream core while retaining conserved N-terminal structural elements.

Together, these analyses identify PF-IIIA as a novel α-synuclein protofilament fold that assembles with canonical PF-IIB2 within an overall MSA Type II–like doublet architecture. The recovery of PF-IIIA-containing filaments from the medial temporal lobe and canonical MSA Type II filaments from the cerebellum demonstrates regional structural heterogeneity within the same individual.

## Discussion

Oligodendroglial cytoplasmic inclusions define MSA, but neuronal α-synuclein inclusions also contribute to its pathological diversity^1,11,12^. Both can coexist in the striatum and brainstem^11,12^, whereas the medial temporal lobe can show prominent neuronal involvement^15,16^. Whether neuronal and oligodendroglial inclusions contain the same α-synuclein folds remains unknown^4,10^. Comparing regions with different inclusion patterns provides an opportunity to examine how filament structure relates to distinct cytopathologies.

In this study, we compared filaments from the medial temporal lobe and cerebellum of an individual with neuropathologically confirmed MSA. We identified a previously undescribed protofilament fold, PF-IIIA, in the medial temporal lobe, whereas cerebellar filaments adopted canonical MSA Type II. This finding expands the known spectrum of MSA-associated folds and demonstrates that distinct PF-A conformations can coexist within a single brain. Because both regions came from the same individual, the differences cannot be explained by genetic or clinical differences between patients. The sampled regions differed in their inclusion types. Oligodendroglial inclusions were widespread, whereas prominent neuronal cytoplasmic inclusions were present in the hippocampus and amygdala but absent from the examined cerebellar sections. The identification of PF-IIIA in the medial temporal lobe and canonical MSA Type II filaments in the cerebellum raises the possibility that PF-IIIA is associated with neuronal inclusions, although the medial temporal tissue contained both inclusion types. This regional correspondence supports investigating whether neurons and oligodendrocytes preferentially accumulate different folds.

Atypical MSA or frontotemporal lobar degeneration with α-synuclein inclusions (FTLD- synuclein) can exhibit abundant eosinophilic, Pick body-like neuronal inclusions in the hippocampus and dentate gyrus, often accompanied by frontotemporal atrophy^4,16^. These cases highlight neuronal α-synuclein accumulation as a substantial component of MSA pathology. Hippocampal α-synuclein pathology also occurs in conventional MSA, generally at more advanced pathological stages and with a lower inclusion burden than in atypical MSA. Brettschneider and colleagues reported hippocampal involvement in phase 3 of 4 in MSA with Parkinsonism (MSA- P)^17^ and phase 4 of 4 in MSA with predominant cerebellar ataxia (MSA-C)^18^. Miki et al. demonstrated that hippocampal and parahippocampal neuronal inclusion burden correlates with memory impairment^19^, whereas Ando et al. emphasized that inclusion density and morphology distinguish atypical MSA from conventional MSA with dementia^15^. Whether these pathological differences correspond to reproducible filament conformations remains unclear.

Previous cryo-EM studies of atypical MSA illustrate the complexity of linking neuronal pathology to a particular fold. Our earlier study identified a Lewy–MSA hybrid fold in the medial temporal lobe of an atypical MSA case^4^, whereas Cullinane and colleagues reported canonical MSA Type II filaments in a case with a similar phenotype^9^. These findings suggest that similar pathological phenotypes can accompany different filament structures, potentially reflecting case variation, regional sampling, or both. Matching structural analysis to histologically examined tissue is important given the marked anatomical asymmetry of atypical MSA. Unlike the study by Cullinane and colleagues, which used different hemispheres for structural and histological analyses^9^, our earlier study examined tissue from the same hemisphere, allowing a closer correlation of filament structure with neuronal pathology^4^. In the present study, we further strengthened this correspondence by preparing frozen sections from the same samples used for cryo-EM and confirmed neuronal α-synuclein-positive inclusions in the sampled medial temporal tissue.

At the molecular level, the novelty of PF-IIIA lies within the protofilament core. Compared with canonical MSA folds, PF-IIIA exhibits substantial structural rearrangements in the C-terminal half of PF-A. APD analysis reveals corresponding differences in side-chain orientations and packing interactions Pronounced PF-A remodeling occurs without a corresponding change in the PF-B fold, demonstrating that a canonical protofilament can partner with different PF-A conformations. The preserved doublet organization and inter-protofilament contacts suggest that alternative PF-A packing arrangements remain compatible with a shared PF-B partner. Thus, substantial changes within one protofilament can be accommodated while key features of the assembled filament remain conserved. Together with previously described chimeric protofilaments and our earlier identification of a Lewy–MSA hybrid fold^4,8^, PF-IIIA illustrates how conserved structural elements can coexist with divergent α-synuclein assemblies.

The coexistence of different PF-A folds raises the question of how distinct conformations arise in different regions of one brain. Local conditions in neurons, oligodendrocytes, or surrounding tissue may favor particular folds^20^. Alternatively, several conformations may form or reach both regions, but differences in growth, stability, retention, or clearance may drive preferential accumulation. Kim et al. reported distinct α-synuclein seeding characteristics in brain regions differing in neuronal and oligodendroglial pathology^21^. This suggests an influence of cellular context on α- synuclein properties but does not establish which folds account for the observed seeding behavior. PF-IIIA in tissue containing prominent neuronal inclusions therefore cannot yet be attributed specifically to inclusion type rather than anatomical location.

Mirroring our findings for α-synuclein filaments in MSA, the coexistence of structurally distinct prion protein aggregates within a single brain has been reported in Creutzfeldt-Jakob disease^22^. α- Synuclein aggregates are proposed to propagate between brain cells via a ‘prion-like’ mechanism^23^. Our discovery of a common PF-IIB2 but brain region-specific PF-A structures raises the possibility that seeding in MSA may be protofilament-specific. Templated addition of α-synuclein monomers to a growing fibril may first occur at the PF-B site, yielding structural fidelity. Seeding at the PF-A site may be a secondary process influenced by cell type-specific cofactors, leading to ‘deformed templating’ and structural divergence^24^. Notably, the short protofilament of the Lewy-MSA hybrid filament we previously identified in atypical MSA also adopts the PF-IIB fold, despite pairing with a structurally distinct Lewy-MSA hybrid protofilament^4^. Intriguingly, seeding with brain-derived MSA type II α-synuclein filaments produced single-protofilament filaments adopting PF-IIB conformations across several cell culture models, without recapitulating the original doublet structures^25^.

A filament structure from one brain region may not represent all α-synuclein folds present in an individual. Neuropathology-guided sampling of multiple regions could reveal structural differences missed by single-region analysis, particularly where neuronal pathology varies. Establishing the extent of variation within individual brains may also help interpret structural differences reported between cases. Such comparisons require sufficient filaments from limited tissue. Using graphene oxide–coated grids^26^, we obtained high-resolution structures from filaments extracted from 1–2 g of tissue per region, facilitating analyses of anatomically restricted areas.

In summary, our study identifies PF-IIIA as a previously undescribed α-synuclein protofilament fold coexisting with canonical MSA Type II filaments in different regions of the same brain. The presence of PF-IIIA in the medial temporal lobe, where neuronal inclusions were prominent, suggests a possible relationship between filament conformation and inclusion type. The conservation of PF-IIB₂ despite divergent PF-A structures also raises the possibility of protofilament-specific templating during α-synuclein propagation. Together, these findings extend the structural diversity of MSA beyond variation between individuals and highlight the importance of regional and cellular pathology in understanding the molecular diversity of α-synuclein assemblies.

## Strengths and Limitations of the study

A major strength of this study is the comparison of high-resolution α-synuclein filament structures from two neuropathologically characterized regions of the same individual, revealing regional structural heterogeneity against a shared genetic and clinical background. Although the regional correspondence between PF-IIIA and neuronal pathology suggests a neuronal association, its cellular origin remains to be established directly. This single-case study cannot determine the prevalence of PF-IIIA or its relationship to clinical phenotype.

## Acknowledgement

We thank the family of the patient for arranging the brain donation. This study was supported by the Rossy Family Foundation, Aligning Science Across Parkinsons (ASAP), the Weston Family Foundation, and the New Frontiers in Research Fund. Cryo-EM data were collected at the Toronto High-Resolution High-Throughput cryo-EM facility, supported by the Canada Foundation for Innovation (CFI) and Ontario Research Fund (ORF). We also thank the infrastructure support provided by the Talos L120C cryo-electron microscope, supported by ORF, which was used for screening cryo-EM samples.

## Author contributions

C.S. and S.E.K. identified the patient, collected clinical information. S.L.F and G.G.K. performed neuropathology. E.R. performed genetic analysis. I.M-V. prepared α-synuclein filament samples.

X.X. and M.E. performed cryo-EM data acquisition, cryo-EM structure determination and structural analyses. M.E. and G.G.K. led the project. X.X., S.L.F., I.M-V., S.E.K., J.C.W., G.G.K., and M.E. contributed to writing the manuscript.

## STAR Methods

### Key Resources Table (to be prepared separately from the main text) Resource Availability

#### Lead contact

Further information and requests for resources and reagents should be directed to and will be fulfilled by the lead contacts,

Masahiro Enomoto Gabor G. Kovacs

## Materials availability

This study did not generate new unique reagents.

## Data availability

The cryo-EM maps from this study have been deposited in the Electron Microscopy Data Bank (EMDB) under accession numbers EMD-79867 for medial temporal filaments and EMD-79774 for cerebellar filaments, respectively. The corresponding refined atomic models have been deposited in the Protein Data Bank (PDB) under accession numbers 39XG for medial temporal filaments and 39TA for cerebellar filaments, respectively.

## Experimental model and study participant details Human samples

The examined specimen was a human brain obtained from the autopsy of a deceased 65-year-old male patient. Consent for autopsy, limited to “brain and neurological tissues” for the purpose of “brain tissue donation for research purposes”, was signed by the spouse of the decedent.

## Methods Details Data reporting

No statistical methods were used to predetermine sample size. The experiments were not randomized, and the investigators were not blinded to allocation during experiments and outcome assessment.

## Neuropathology

Brain tissue was examined in the frame of an ongoing Brain Donation program of the Toronto Western Hospital following the approval of the local ethics committee (University Health Network Research Ethics Board, Nr. 20-5258) and adherence to appropriate consent protocols.

Formalin-fixed, 4.5 μm paraffin-embedded tissue sections comprising the frontal, temporal, parietal, anterior cingulate, motor and occipital cortices, anterior and posterior regions of the basal ganglia, thalamus, midbrain, pons, medulla oblongata and cerebellum with dentate nucleus were stained with hematoxylin, eosin, and luxol fast blue (HE/LFB), and Gallyas Braak silver staining. For the neuropathology analysis including the evaluation for mixed pathology, various immunostains including p-tau (clone AT8, 1:1000, Invitrogen/ThermoFisher, Carlsbad, USA), Aβ (Clone 6F/3D, 1:50, Dako, Glostrup, Denmark), a-synuclein (clone 5G4, 1:4000, Analytikjena, Jena, Germany)^27^, phosphorylated TDP-43 (clone 11-9, 1:2000, CosmoBio, Tokyo, Japan) and anti-p62 (3/P62 LCK LIGAND, 1:500 BD Company, NJ, United States) were employed. Alzheimer’s Disease (AD)-related pathology was assessed through the Braak neurofibrillary tangle (NFT) stage^28^ and Thal phase for Aβ deposition^29^.

We also sampled frozen brain tissue corresponding to the sample that was processed for cryo-EM. The frozen sections were fixed in 95% ethanol for 30 seconds, rinsed briefly in running tap water, and stained with Harris haematoxylin for 1 min. Sections were then rinsed, dehydrated through graded alcohols (70–100%, 4–6 dips each), cleared in 2–3 changes of xylene (3–6 dips each), and coverslipped with Permount. Sections were subsequently immunostained for phosphorylated α- synuclein (P-Syn antibody is EY1536Y, from Abcam, 1:500, rabbit monoclonal antibodyand p62 (clone 3/p62 LCK ligand, 1:500; BD Biosciences, NJ, USA).

## Genetic analysis

Genomic DNA was isolated from brain using a QIAGEN kit. The entire open reading frame of *SNCA* was Sanger sequenced. Gene dosage of the *SNCA* locus on chromosome 4 was determined by inspecting the B allele frequency and LogR Ratio plots from Neurobooster array data (155 variants) using GenomeStudio (Illumina Inc., CA).

## Protein extraction

Detergent-extracted brain homogenates [10% (w/v)] were prepared in PBS using a gentle-MACS Octo Dissociator (Miltenyi BioTec). Nine volumes of brain homogenate were then mixed with one volume of 10× detergent buffer [5% (v/v) Nonidet P-40, 5% (w/v) sodium deoxycholate in PBS] containing Pierce Universal Nuclease (ThermoFisher) and Halt Phosphatase Inhibitor (ThermoFisher), and then incubated on ice for 20 min. Samples were clarified by centrifugation at 1000x *g* for 5 min at 4 °C to generate detergent-extracted brain homogenate, as previously described.^30^ The supernatant was collected and aliquoted in 0.5-ml low protein binding tubes (Eppendorf) to avoid excessive freeze–thaw cycles. A bicinchoninic acid protein (BCA) assay (ThermoFisher) was performed to determine the total protein concentration of all samples.

The sarkosyl-insoluble extraction was performed as previously described, with minor modifications^31^. Briefly, 1 gram of tissue was homogenised in 20% (v/w) extraction buffer consisting of 10 mM Tris-HCl, pH 7.5, 0.8 M NaCl, 10% sucrose and 1 mM EGTA (Sigma). Homogenates were brought to 2% sarkosyl (Sigma) and incubated for 30 min. at 37 °C. Following a 13 min. centrifugation at 10,000 g, the supernatants were spun at 100,000 g for 30 min. The pellets were resuspended in 500 μl/g extraction buffer and centrifuged at 3,000 g for 5 min. The supernatants were diluted 3-fold in 50 mM Tris-HCl, pH 7.5, containing 0.15 M NaCl, 10% sucrose and 0.2% sarkosyl (Sigma), and spun at 166,000 g for 40 min. Sarkosyl-insoluble pellets were resuspended in 100 μl/g of PBS (Invitrogen).

## Electron cryo-microscopy

The sarkosyl-insoluble extracts were applied to homemade glow-discharged holey gold/carbon grids^32^ coated with homemade graphene oxide flakes ^26,33^ and plunge-frozen in liquid ethane using a Thermo Fisher Vitrobot Mark IV. High-resolution data collection was then performed with a Titan Krios G3 electron microscope operating at 300 kV and equipped with a Falcon 4i camera and a Selectris X energy filter (Thermo Fisher Scientific) using a slit width of 10 eV. Electron microscopy data acquisition was automated with the EPU software (Thermo Fisher Scientific).

Movies were acquired in electron event representation mode^34^ at 130,000× nominal magnification, corresponding to a calibrated pixel size of 0.93 Å, with a total exposure of ∼40 e-/Å2, and subsequently divided into 40 exposure fractions. Further details are provided in **Supplementary Table S1**.

## Helical reconstruction

Movie frames were corrected for beam-induced motion and dose-weighted using the motion correction implementation in RELION 5.0 ^35^. The aligned, non-dose-weighted micrographs were used to estimate the contrast transfer function (CTF) with CTFFIND4 v4.1^36^. All subsequent image processing was performed using the helical reconstruction pipeline in RELION 5.0. For both datasets, α-synuclein filaments were manually picked. Segments spanning an entire helical crossover were extracted for reference-free 2D classification using an inter-box distance of 14.1 Å. For the medial temporal lobe dataset, segments were initially extracted with a box size of 1,400 pixels at a pixel size of 0.93 Å and downsampled to 350 pixels. Reference-free 2D classification was performed to remove segments contributing to suboptimal class averages. The remaining segments were then re-extracted with a box size of 768 pixels at the original pixel size (0.93 Å), downsampled to 256 pixels, and subjected to a second round of reference-free 2D classification to remove additional suboptimal segments. The remaining segments were subsequently re-extracted from the original micrographs without downsampling using a box size of 354 pixels and subjected to further reference-free 2D classification. For the cerebellum dataset, segments were initially extracted with a box size of 900 pixels at a pixel size of 0.93 Å and downsampled to 300 pixels. Reference-free 2D classification was performed to remove segments contributing to suboptimal class averages. The remaining segments were then re-extracted from the original micrographs without downsampling using a box size of 354 pixels and subjected to further reference-free 2D classification. For both datasets, the resulting high-quality 2D class averages were used to generate an initial *de novo* 3D reference model with the relion_helix_inimodel2d program^37^. An initial 3D auto- refinement was then performed using the re-extracted segments and the *de novo* reference model low-pass filtered to 15 Å. Map quality and resolution were further improved through iterative rounds of 3D classification and 3D auto-refinement. Bayesian polishing and CTF refinement were subsequently performed to further improve the resolution of the reconstructions. The final reconstructions were sharpened using the standard post-processing procedures implemented in RELION 5.0. Overall resolution was estimated from the gold-standard Fourier shell correlation (FSC) at the 0.143 criterion between two independently refined half-maps, with phase randomization used to correct for convolution effects introduced by a generous soft-edged solvent mask. Initial atomic models were built manually in ISOLDE^38^ and Coot^39^. Atomic coordinates were refined using Phenix^40^. The final models were validated with MolProbity^41^. Figures were prepared with UCSF ChimeraX^42^. Model statistics are summarized in **Supplementary Table S1**.

## Amyloid packing difference (APD) analysis

APD analysis was performed as described previously^13^. Pairwise XY-APD values were calculated between MT-filament, Cere-filament, and previously reported α-synuclein protofilament structures using the original APD scripts^13^. Hierarchical clustering of the pairwise XY-APD matrix was performed in Python using the SciPy library, and dendrograms were generated with Matplotlib. APD schematic representations were generated using the original APD scripts.

## References

1. Stefanova, N., and Wenning, G.K. (2023). Multiple system atrophy: at the crossroads of cellular, molecular and genetic mechanisms. Nat Rev Neurosci 24, 334–346. 10.1038/s41583-023-00697-7.

2. Goedert, M., Jakes, R., and Spillantini, M.G. (2017). The Synucleinopathies: Twenty Years On. J Parkinsons Dis 7, S51–S69. 10.3233/JPD-179005.

3. Spillantini, M.G., and Goedert, M. (2000). The alpha-synucleinopathies: Parkinson’s disease, dementia with Lewy bodies, and multiple system atrophy. Ann N Y Acad Sci 920, 16–27. 10.1111/j.1749-6632.2000.tb06900.x.

4. Enomoto, M., Martinez-Valbuena, I., Forrest, S.L., Xu, X., Munhoz, R.P., Li, J., Rogaeva, E., Lang, A.E., and Kovacs, G.G. (2025). Lewy-MSA hybrid fold drives distinct neuronal α-synuclein pathology. Communications Biology 8, 929.

5. do Carmo Goncalves, P., Fernandez, I., and Fernandez Outon, C.O. (2026). Structural biology of alpha-synuclein amyloid filaments: Implications in synucleinopathies. J Parkinsons Dis 16, 1094–1114. 10.1177/1877718X261461375.

6. Schweighauser, M., Shi, Y., Tarutani, A., Kametani, F., Murzin, A.G., Ghetti, B., Matsubara, T., Tomita, T., Ando, T., and Hasegawa, K. (2020). Structures of α-synuclein filaments from multiple system atrophy. Nature 585, 464–469.

7. Yang, Y., Shi, Y., Schweighauser, M., Zhang, X., Kotecha, A., Murzin, A.G., Garringer, H.J., Cullinane, P.W., Saito, Y., and Foroud, T. (2022). Structures of α-synuclein filaments from human brains with Lewy pathology. Nature 610, 791–795.

8. Yan, N.L., Candido, F., Tse, E., Melo, A.A., Prusiner, S.B., Mordes, D.A., Southworth, D.R., Paras, N.A., and Merz, G.E. (2025). Cryo-EM structure of a novel α-synuclein filament subtype from multiple system atrophy. Febs Letters 599, 33–40.

9. Cullinane, P.W., Yang, Y., Chelban, V., Goh, Y.Y., Ebanks, K., Curless, T., Wrigley, S., de Pablo-Fernandez, E., Holton, J., Peak-Chew, S., et al. (2025). Identical Seeding Characteristics and Cryo-EM Filament Structures in FTLD-Synuclein and Typical Multiple System Atrophy. Neuropathol Appl Neurobiol 51, e70013. 10.1111/nan.70013.

10. Boing, C., Di Fabrizio, M., Burger, D., Bol, J., Huisman, E., Rozemuller, A.J.M., van de Berg, W.D.J., Stahlberg, H., and Lewis, A.J. (2024). Distinct ultrastructural phenotypes of glial and neuronal alpha-synuclein inclusions in multiple system atrophy. Brain 147, 3727–3741. 10.1093/brain/awae137.

11. Dhillon, J.S., Trejo-Lopez, J.A., Riffe, C., McFarland, N.R., Hiser, W.M., Giasson, B.I., and Yachnis, A.T. (2019). Dissecting alpha-synuclein inclusion pathology diversity in multiple system atrophy: implications for the prion-like transmission hypothesis. Lab Invest 99, 982–992. 10.1038/s41374-019-0198-9.

12. Hass, E.W., Sorrentino, Z.A., Lloyd, G.M., McFarland, N.R., Prokop, S., and Giasson, B.I. (2021). Robust alpha-synuclein pathology in select brainstem neuronal populations is a potential instigator of multiple system atrophy. Acta Neuropathol Commun 9, 80. 10.1186/s40478-021-01173-y.

13. Scheres, S.H.W. (2026). The amyloid packing difference: A pairwise comparison metric for amyloid structures. Structure 34, 1061–1071 e1061. 10.1016/j.str.2026.04.014.

14. Yang, Y., Garringer, H.J., Shi, Y., Lövestam, S., Peak-Chew, S., Zhang, X., Kotecha, A., Bacioglu, M., Koto, A., and Takao, M. (2023). New SNCA mutation and structures of α- synuclein filaments from juvenile-onset synucleinopathy. Acta Neuropathologica 145, 561–572.

15. Ando, T., Riku, Y., Akagi, A., Miyahara, H., Hirano, M., Ikeda, T., Yabata, H., Koizumi, R., Oba, C., Morozumi, S., et al. (2022). Multiple system atrophy variant with severe hippocampal pathology. Brain Pathol 32, e13002. 10.1111/bpa.13002.

16. Aoki, N., Boyer, P.J., Lund, C., Lin, W.L., Koga, S., Ross, O.A., Weiner, M., Lipton, A., Powers, J.M., White, C.L., 3rd, and Dickson, D.W. (2015). Atypical multiple system atrophy is a new subtype of frontotemporal lobar degeneration: frontotemporal lobar degeneration associated with alpha-synuclein. Acta Neuropathol 130, 93–105. 10.1007/s00401-015-1442-z.

17. Brettschneider, J., Suh, E., Robinson, J.L., Fang, L., Lee, E.B., Irwin, D.J., Grossman, M., Van Deerlin, V.M., Lee, V.M., and Trojanowski, J.Q. (2018). Converging Patterns of alpha-Synuclein Pathology in Multiple System Atrophy. J Neuropathol Exp Neurol 77, 1005–1016. 10.1093/jnen/nly080.

18. Brettschneider, J., Irwin, D.J., Boluda, S., Byrne, M.D., Fang, L., Lee, E.B., Robinson, J.L., Suh, E., Van Deerlin, V.M., Toledo, J.B., et al. (2017). Progression of alpha-synuclein pathology in multiple system atrophy of the cerebellar type. Neuropathol Appl Neurobiol 43, 315–329. 10.1111/nan.12362.

19. Miki, Y., Foti, S.C., Hansen, D., Strand, K.M., Asi, Y.T., Tsushima, E., Jaunmuktane, Z., Lees, A.J., Warner, T.T., Quinn, N., et al. (2020). Hippocampal alpha-synuclein pathology correlates with memory impairment in multiple system atrophy. Brain 143, 1798–1810. 10.1093/brain/awaa126.

20. Peng, C., Gathagan, R.J., Covell, D.J., Medellin, C., Stieber, A., Robinson, J.L., Zhang, B., Pitkin, R.M., Olufemi, M.F., Luk, K.C., et al. (2018). Cellular milieu imparts distinct pathological alpha-synuclein strains in alpha-synucleinopathies. Nature 557, 558–563. 10.1038/s41586-018-0104-4.

21. Kim, A., Martinez-Valbuena, I., Danics, K., Forrest, S.L., and Kovacs, G.G. (2025). Contribution of alpha-synuclein cytopathologies to distinct seeding of misfolded alpha- synuclein. Brain Pathol 35, e70024. 10.1111/bpa.70024.

22. Cali, I., Castellani, R., Alshekhlee, A., Cohen, Y., Blevins, J., Yuan, J., Langeveld, J.P., Parchi, P., Safar, J.G., Zou, W.Q., and Gambetti, P. (2009). Co-existence of scrapie prion protein types 1 and 2 in sporadic Creutzfeldt-Jakob disease: its effect on the phenotype and prion-type characteristics. Brain 132, 2643–2658. 10.1093/brain/awp196.

23. Steiner, J.A., Quansah, E., and Brundin, P. (2018). The concept of alpha-synuclein as a prion-like protein: ten years after. Cell Tissue Res 373, 161–173. 10.1007/s00441-018-2814-1.

24. Makarava, N., and Baskakov, I.V. (2013). The evolution of transmissible prions: the role of deformed templating. PLoS Pathog 9, e1003759. 10.1371/journal.ppat.1003759.

25. Zhang, D., Matic, M., Tarutani, A., Murzin, A.G., Huang, M., Macdonald, J.A., Katsinelos, T., Cullinane, P.C., Hasegawa, K., Jaunmuktane, Z., et al. (2026). Oligodendrocytes may provide the extra density of α-synuclein filaments from multiple system atrophy. bioRxiv, 2026.2010.2029.762735. 10.64898/2026.10.29.762735.

26. Palovcak, E., Wang, F., Zheng, S.Q., Yu, Z., Li, S., Betegon, M., Bulkley, D., Agard, D.A., and Cheng, Y. (2018). A simple and robust procedure for preparing graphene-oxide cryo- EM grids. J Struct Biol 204, 80–84. 10.1016/j.jsb.2018.07.007.

27. Kovacs, G.G., Wagner, U., Dumont, B., Pikkarainen, M., Osman, A.A., Streichenberger, N., Leisser, I., Verchere, J., Baron, T., Alafuzoff, I., et al. (2012). An antibody with high reactivity for disease-associated alpha-synuclein reveals extensive brain pathology. Acta Neuropathol 124, 37–50. 10.1007/s00401-012-0964-x.

28. Braak, H., Alafuzoff, I., Arzberger, T., Kretzschmar, H., and Del Tredici, K. (2006). Staging of Alzheimer disease-associated neurofibrillary pathology using paraffin sections and immunocytochemistry. Acta Neuropathol 112, 389–404. 10.1007/s00401-006-0127-z.

29. Thal, D.R., Rub, U., Orantes, M., and Braak, H. (2002). Phases of A beta-deposition in the human brain and its relevance for the development of AD. Neurology 58, 1791–1800. 10.1212/wnl.58.12.1791.

30. Lau, A., So, R.W.L., Lau, H.H.C., Sang, J.C., Ruiz-Riquelme, A., Fleck, S.C., Stuart, E., Menon, S., Visanji, N.P., Meisl, G., et al. (2020). alpha-Synuclein strains target distinct brain regions and cell types. Nat Neurosci 23, 21–31. 10.1038/s41593-019-0541-x.

31. Schweighauser, M., Shi, Y., Tarutani, A., Kametani, F., Murzin, A.G., Ghetti, B., Matsubara, T., Tomita, T., Ando, T., Hasegawa, K., et al. (2020). Structures of alpha- synuclein filaments from multiple system atrophy. Nature 585, 464–469. 10.1038/s41586-020-2317-6.

32. Marr, C.R., Benlekbir, S., and Rubinstein, J.L. (2014). Fabrication of carbon films with∼ 500 nm holes for cryo-EM with a direct detector device. Journal of Structural Biology 185, 42–47.

33. Chen, D., Feng, H., and Li, J. (2012). Graphene oxide: preparation, functionalization, and electrochemical applications. Chem Rev 112, 6027–6053. 10.1021/cr300115g.

34. Guo, H., Franken, E., Deng, Y., Benlekbir, S., Singla Lezcano, G., Janssen, B., Yu, L., Ripstein, Z.A., Tan, Y.Z., and Rubinstein, J.L. (2020). Electron-event representation data enable efficient cryoEM file storage with full preservation of spatial and temporal resolution. IUCrJ 7, 860–869.

35. Scheres, S.H. (2012). RELION: implementation of a Bayesian approach to cryo-EM structure determination. J Struct Biol 180, 519–530. 10.1016/j.jsb.2012.09.006.

36. Rohou, A., and Grigorieff, N. (2015). CTFFIND4: Fast and accurate defocus estimation from electron micrographs. J Struct Biol 192, 216–221. 10.1016/j.jsb.2015.08.008.

37. Scheres, S.H.W. (2020). Amyloid structure determination in RELION-3.1. Acta Crystallogr D Struct Biol 76, 94–101. 10.1107/S2059798319016577.

38. Croll, T.I. (2018). ISOLDE: a physically realistic environment for model building into low- resolution electron-density maps. Acta Crystallogr D Struct Biol 74, 519–530. 10.1107/S2059798318002425.

39. Emsley, P., Lohkamp, B., Scott, W.G., and Cowtan, K. (2010). Features and development of Coot. Acta Crystallogr D Biol Crystallogr 66, 486–501. 10.1107/S0907444910007493.

40. Liebschner, D., Afonine, P.V., Baker, M.L., Bunkoczi, G., Chen, V.B., Croll, T.I., Hintze, B., Hung, L.W., Jain, S., McCoy, A.J., et al. (2019). Macromolecular structure determination using X-rays, neutrons and electrons: recent developments in Phenix. Acta Crystallogr D Struct Biol 75, 861–877. 10.1107/S2059798319011471.

41. Williams, C.J., Headd, J.J., Moriarty, N.W., Prisant, M.G., Videau, L.L., Deis, L.N., Verma, V., Keedy, D.A., Hintze, B.J., Chen, V.B., et al. (2018). MolProbity: More and better reference data for improved all-atom structure validation. Protein Sci 27, 293–315. 10.1002/pro.3330.

42. Meng, E.C., Goddard, T.D., Pettersen, E.F., Couch, G.S., Pearson, Z.J., Morris, J.H., and Ferrin, T.E. (2023). UCSF ChimeraX: Tools for structure building and analysis. Protein Sci 32, e4792. 10.1002/pro.4792.

