## Supplemental Figures and Table for "α-Synuclein structural heterogeneity associates with distinct cytopathologies in multiple system atrophy"

#### Supplementary Figure 1

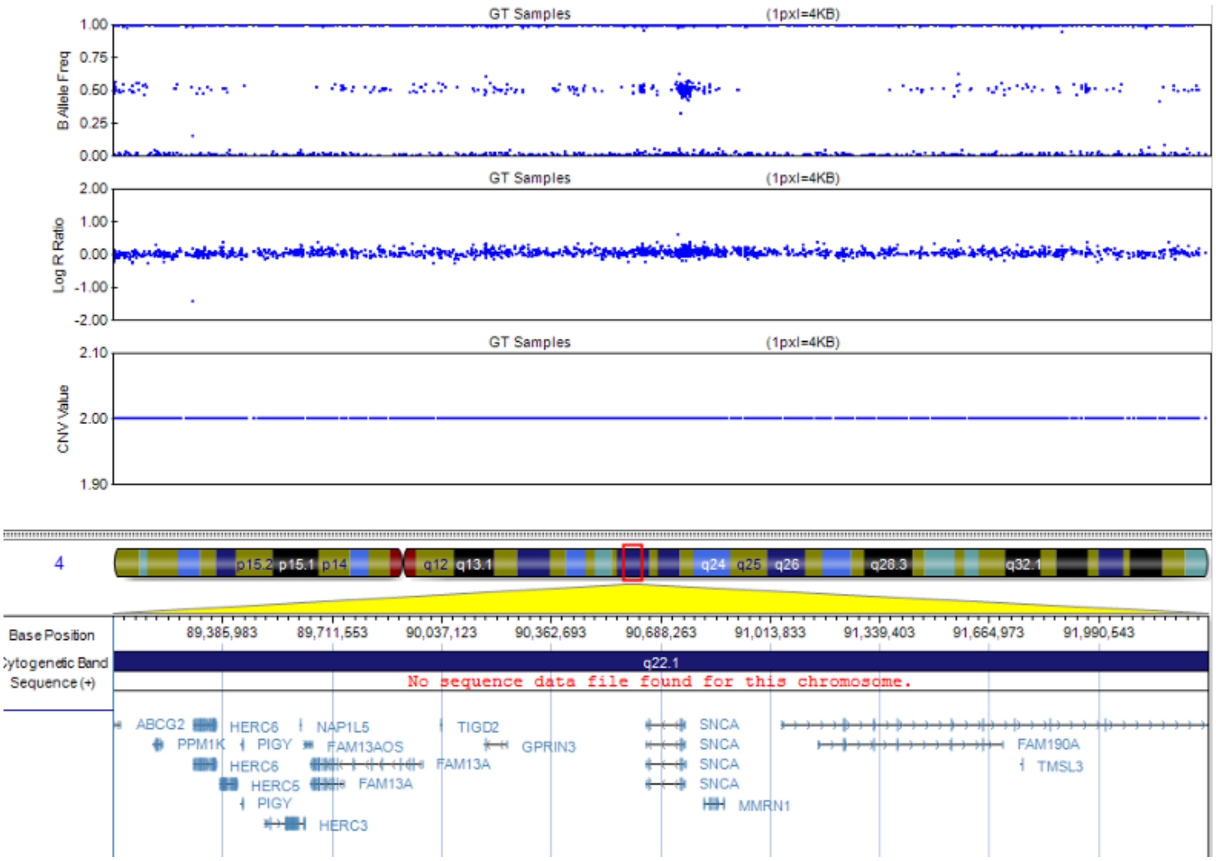

**Supplementary Figure 1. Screenshot of B allele frequency and LogR ratio plots for the MSA case in chromosome browser using GenomeStudio.**

#### Supplementary Figure 2

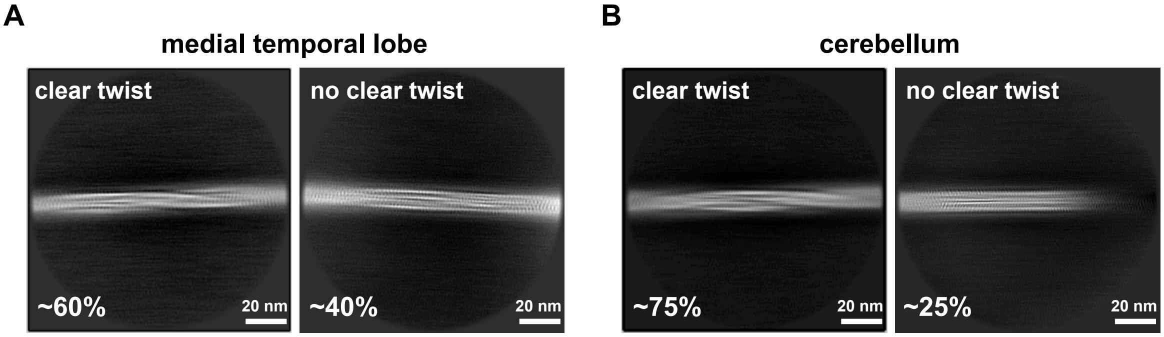

**Supplementary Figure 2. Representative 2D class averages of α-synuclein filament segments.** **(A)** Medial temporal lobe. **(B)** Cerebellum. For each region, representative class averages show segments with a clear twist (left) and no clear twist (right). Percentages indicate the approximate fraction of classified segments in each category. Scale bars, 20 nm.

#### Supplementary Figure 3

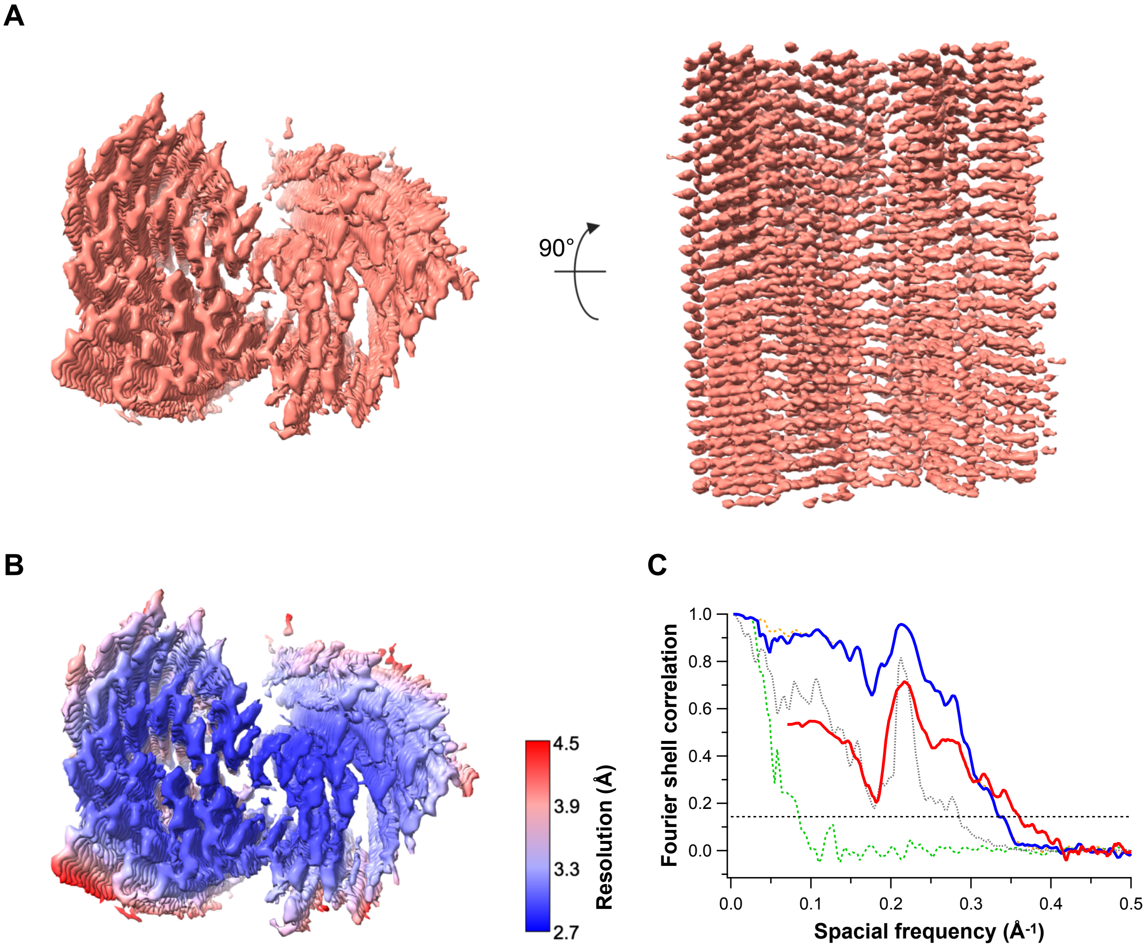

**Supplementary Figure 3. Cryo-EM reconstruction of α-synuclein filaments from the medial temporal lobe.**

**(A)** End and side views of the reconstructed filament density, rotated by 90° relative to each other. **(B)** Cryo-EM map colored by local resolution. **(C)** Fourier shell correlation (FSC) curves from half-map analysis and map–model validation. Half-map FSC curves are shown as corrected (solid blue), unmasked (dotted gray), phase-randomized masked (dotted green), and masked (dotted orange). The masked FSC curve was omitted for clarity because it nearly completely overlapped with the corrected FSC curve. The map–model FSC calculated using Phenix is shown in solid red. The horizontal dotted line indicates the FSC threshold of 0.143, where the corrected half-map FSC yields a global resolution of 3.0 Å.

#### Supplementary Figure 4

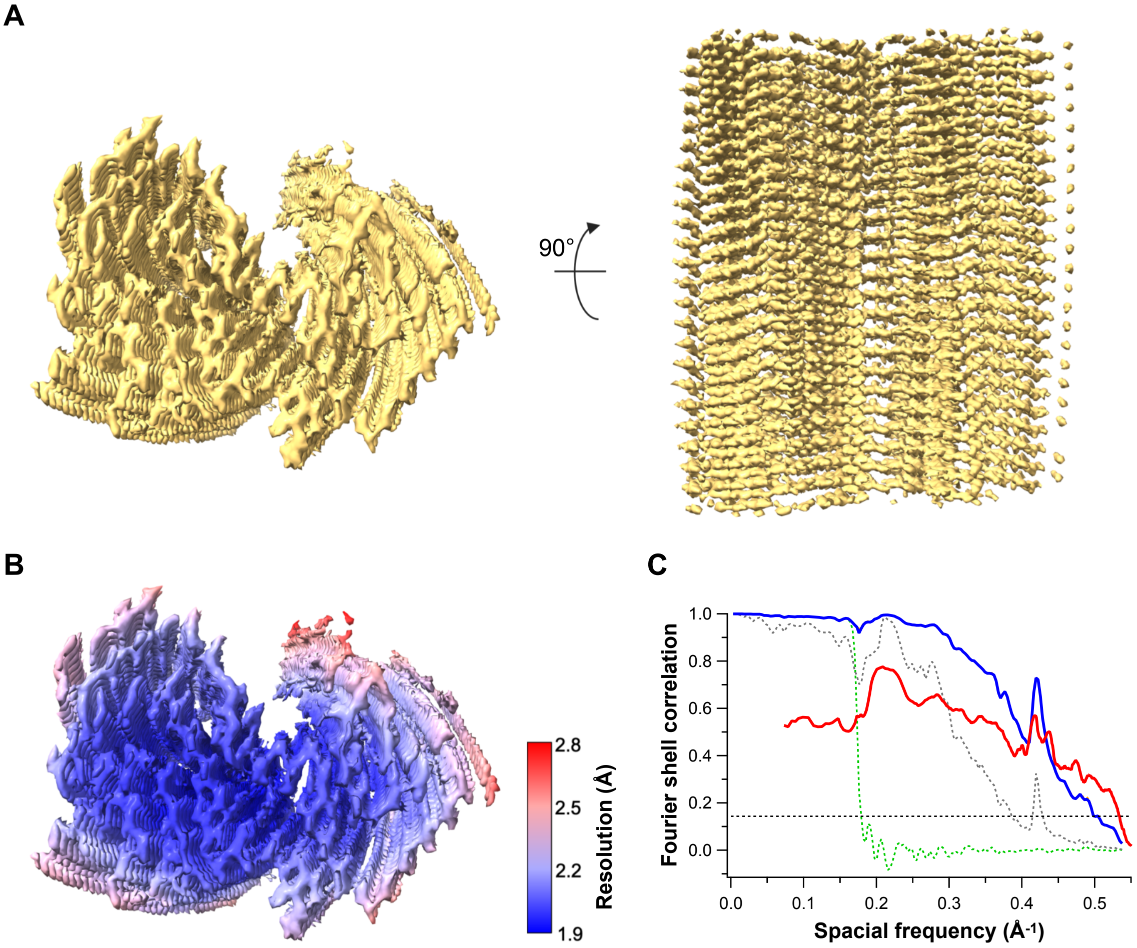

**Supplementary Figure 4. Cryo-EM reconstruction of α-synuclein filaments from the cerebellum.**

**(A)** End and side views of the reconstructed filament density, rotated by 90° relative to each other. **(B)** Cryo-EM map colored by local resolution. **(C)** Fourier shell correlation (FSC) curves from half-map analysis and map–model validation. Half-map FSC curves are shown as corrected (solid blue), unmasked (dotted gray), and phase-randomized masked (dotted green). The masked FSC curve was omitted for clarity because it nearly completely overlapped with the corrected FSC curve. The map–model FSC calculated using Phenix is shown in solid red. The horizontal dotted line indicates the FSC threshold of 0.143, where the corrected half-map FSC yields a global resolution of 2.0 Å.

### Supplementary Figure 5

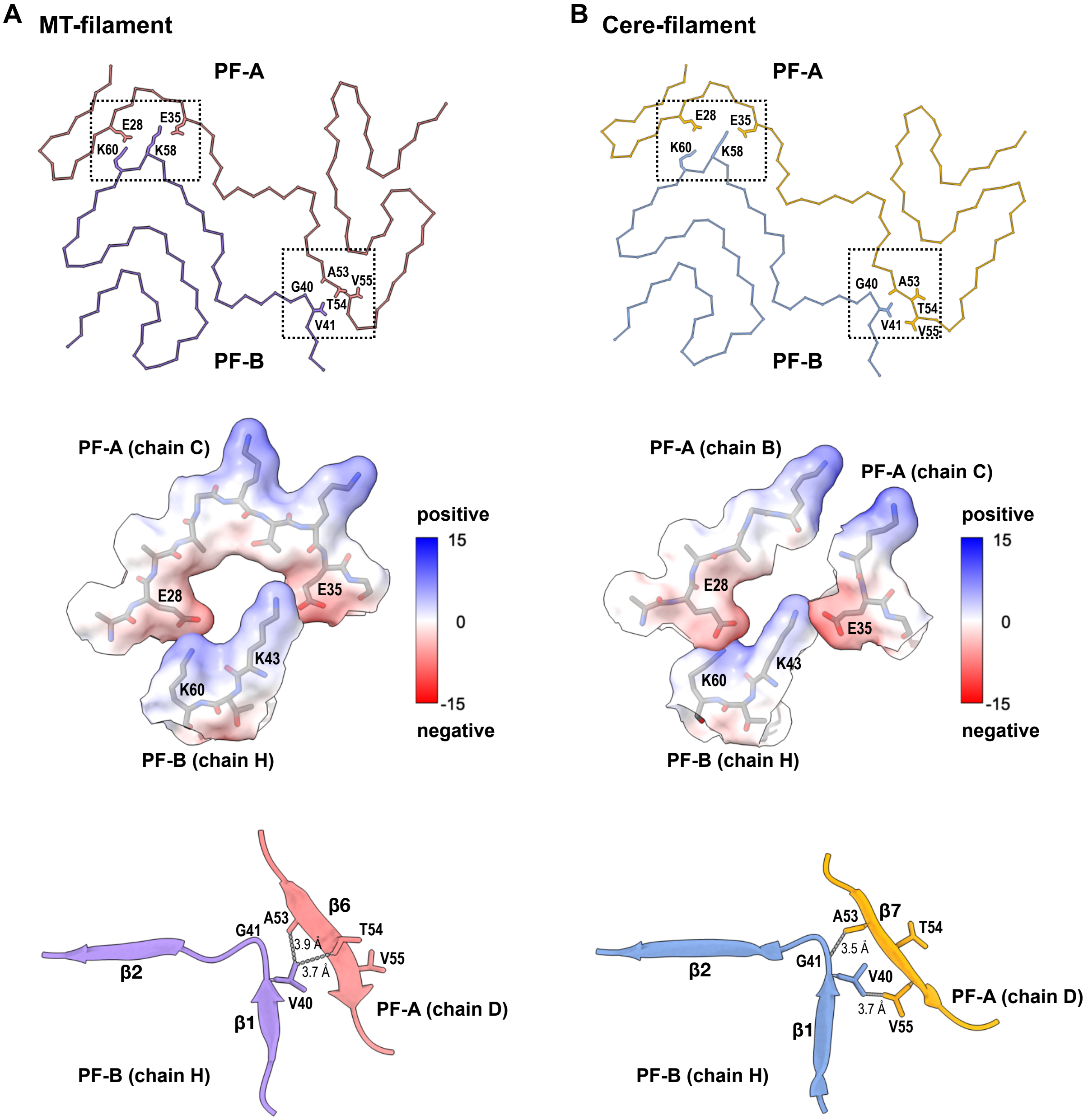

**Supplementary Figure 5. Comparison of inter-protofilament interactions in medial temporal and cerebellar α-synuclein filaments.**
**(A, B)** Inter-protofilament interfaces of the medial temporal (MT-, **A**) and cerebellar (Cere-, **B**) filaments. **Top:** schematic representations showing PF-A (chain C) and PF-B (chain H) for simplicity. Dashed boxes indicate regions involved in electrostatic interactions and close packing between protofilaments. **Middle:** enlarged electrostatic surface representations of the charged interfaces, colored according to Coulombic potential from −15 (red, negative) to +15 (blue, positive) kcal/(mol·e). In the MT-filament, E28 and E35 are shown from the same axial rung of PF-A (chain C), whereas in the Cere-filament, E28 and E35 are from different axial rungs of PF-A (chains B and C, respectively) interact with K60 and K58 of PF-B (chain H). **Bottom:** enlarged views showing close packing between a β-strand of PF-A (β6 in the MT-filament and β7 in the Cere-filament; chain D in both structures) and the β1–β2 loop of PF-B (chain H). Thus, the packing interactions shown in the bottom panels involve a different axial rung of PF-A from that depicted in the simplified top panels.

#### Supplementary Figure 6

**PF-IIA (Cere-PF-A)**

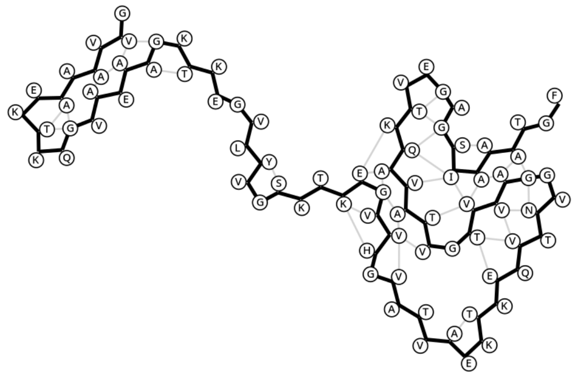

**PF-IIA (6XYQ)**

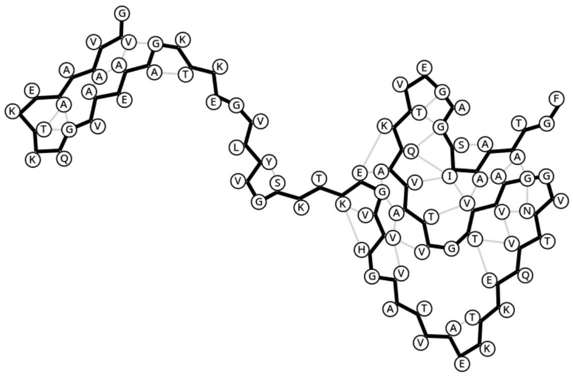

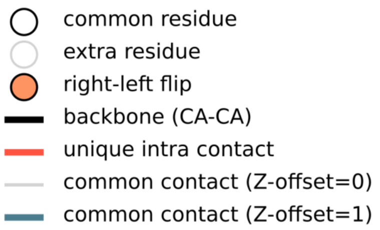

| **Cere-PF-IIA vs PF-IIA (6XYQ)** | |
|---|---|
| residues in Cere-PF-IIA | 81 |
| residues in PF-IIA (6XYQ) | 81 |
| residues in common | 81 |
| extra residues in Cere-PFII-A | 0 |
| **For common residues** | |
| right-left flips | 0 |
| unique intra-contacts | 0 |
| Z-offsets | 0 |
| XY-differences | 0 |
| XYZ-differences | 0 |
| **PF-IIIA APD (XY)** | **0%** |
| **PF-IIIA APD (XYZ)** | **0%** |

**Supplementary Figure 6. Conserved packing interactions in Cere-PF-A and canonical PF-IIA.**
APD schematic comparison of Cere-PF-A (chain C; left) and canonical PF-IIA (PDB 6XYQ, chain A; right). Circles represent residues labeled using single-letter amino acid codes; black lines connect consecutive Cα atoms, and gray lines indicate shared packing contacts with a Z-offset of 0. Both structures contain 81 ordered residues, with no differences in residue orientations, packing contacts, or axial offsets assessed by APD, yielding XY-APD and XYZ-APD values of 0%.

**Supplementary Table 1. Statistics for cryo-EM data acquisition and atomic model building**

| | **MT-filament**<br>EMD-79867, PDB ID 39XG | **Cere-filament**<br>EMD-79774, PDB ID 39TA |
|---|---|---|
| **Data acquisition** | | |
| Microscope | Titan Krios G3 | Titan Krios G3 |
| Detector | Falcon 4i | Falcon 4i |
| Energy filter slit (eV) | 10 | 10 |
| Magnification | 130,000 | 130,000 |
| Voltage (kV) | 300 | 300 |
| Exposure rate (e/pix/s) | 7.6 | 7.5 |
| Electron dose (e/Å$^2$) | 40 | 40 |
| Defocus range (μm) | 0.8-2.0 | 0.8-2.0 |
| Pixel size (Å) | 0.93 | 0.93 |
| **Map refinement** | | |
| Symmetry imposed | C1 | C1 |
| Final particle images (No.) | 10.805 | 106,479 |
| Map resolution (Å) | 3.0 | 2.0 |
| FSC threshold | 0.143 | 0.143 |
| Helical twist (°) | -1.28 | -1.38 |
| Helical rise (Å) | 4.73 | 4.75 |
| **Model refinement** | | |
| Model resolution (Å) | 2.9 | 1.9 |
| FSC threshold | 0.143 | 0.143 |
| Map sharpening B factor (Å$^2$) | -47.1268 | -24.1496 |
| Model composition | | |
| Non-hydrogen atoms | 5.175 | 4,955 |
| Protein residues | 750 | 725 |
| B factor (Å$^2$) | | |
| Protein | 36.50 | 31.72 |
| R.m.s deviation | | |
| Bond lengths (Å) | 0.006 | 0.007 |
| Bond angles (°) | 2.947 | 1.293 |
| Validation | | |
| MolProbity score | 2.10 | 1.75 |
| Clashscore | 11.71 | 5.73 |
| Rotamer outliers (%) | 0.00 | 0.00 |
| Ramachandran plot | | |
| Favored (%) | 91.10 | 93.19 |
| Allowed (%) | 8.77 | 6.81 |
| Outliers (%) | 0.14 | 0.00 |
